# Recombination rate inference via deep learning is limited by sequence diversity

**DOI:** 10.1101/2022.07.01.498489

**Authors:** Mackenzie M. Johnson, Claus O. Wilke

## Abstract

A common inference task in population genetics is to estimate recombination rate from multiple sequence alignments. Traditionally, recombination rate estimators have been developed from biologically-informed, statistical models, but more recently deep learning models have been employed for this task. While deep learning approaches offer unique advantages, their performance is inconsistent across the range of potential recombination rates. Here, we generate and characterize data sets (genotype alignments with known recombination rates) for use by deep learning estimators and assess how their features limit estimator performance. We find that certain input parameter regimes produce genotype alignments with low sequence diversity, which are inherently information-limited. We next test how estimator performance is impacted by training and evaluating neural networks on data sets with varying degrees of diversity. The inclusion of genotype alignments with low diversity at high frequency results in considerable performance declines across two different network architectures. In aggregate, our results suggest that genotype alignments have inherent information limits when sequence diversity is low, and these limitations need to be considered both when training deep learning recombination rate estimators and when using them in inference applications.

## Introduction

Recombination is a major evolutionary force introducing diversity into eukaryotic genomes. During sexual reproduction, maternally- and paternally-inherited chromosomes line up and recombine, exchanging homologous segments of DNA via crossover events. The byproduct of this process is the creation of novel haplotypes, combinations of alleles found on the same chromosome, on which selection can act. A major focus of population genetics has been to quantify the rate of recombination to better understand the historical effects of recombination in a population (Hahn, 2019). Directly measuring recombination rate in natural and lab populations is often infeasible or impractical. Thus, population recombination rates are commonly inferred using statistical estimators applied to large, high-throughput sequencing data sets (Hahn, 2019; Peñalba and Wolf, 2020). Understanding the historical pattern of recombination in individual regions and across the genome is crucial; these estimates provide information necessary to detect genome wide associations, changes in demography (population growth and bottlenecks), and events of recent positive selection (Pritchard and Przeworski, 2001; Ardlie et al., 2002; Sodeland et al., 2011; Sabeti et al., 2002).

Recombination rate is known to vary widely across genomes and between individuals, sexes, populations, and species (Stapley et al., 2017; Sardell and Kirkpatrick, 2020; Peñalba and Wolf, 2020). However, the causes of and extent of this observed variation remain under-explored as researchers are limited by the tools available (Peñalba and Wolf, 2020; Adrion et al., 2020). State-of-the-art estimators of population recombination rate include composite likelihood methods (McVean et al., 2002; Stumpf and McVean, 2003; Chan et al., 2012) and supervised machine learning methods (Lin et al., 2013; Gao et al., 2016). To achieve accurate estimates for a given population, these methods require considerable computational resources and large, high-quality alignments of phased haplotypes. Additionally, these methods rely on existing population genetic theory to connect features of the population data to the underlying evolutionary process that generated the observed haplotypes (Flagel et al., 2019). Recent studies have found that some common pitfalls of conventional methods can be avoided by training deep neural networks with simulation data (Chan et al., 2018; Flagel et al., 2019; Adrion et al., 2020). Neural network estimators are particularly promising because of their applicability to populations where conventional methods cannot be used to estimate recombination rate (Flagel et al., 2019). This methodological advancement has the potential to expand our understanding of how and where recombination rate varies, elucidating the role of recombination in evolutionary history. Yet, the performance advantage of neural networks (respective to conventional methods) appears to be dependent on the recombination rate; neural networks work best when recombination rates are relatively high, but are outperformed by traditional estimators when recombination rate is low (Flagel et al., 2019). The conditions under which neural networks perform poorly for recombination rate estimation are not fully understood.

Here, we assess limitations of deep learning models trained for population recombination rate estimation by: 1) simulating alignments under different parameter regimes and quantifying their sequence diversity (information quality) and 2) analyzing the impact of low-diversity alignments on the performance of convolutional neural networks (CNNs) trained as estimators (LeCun et al., 1998). Each coalescent simulation produces a genotype alignment (a matrix of 0s and 1s) and a vector of positional information (chromosomal coordinates) for all variable sites in the population. We generate data sets from a range of input parameter values, selected to either maximize or minimize information, and compare the variation across genotype alignments within a given regime. Parameter regimes with low sequence diversity are expected to have a detrimental impact on network training. We demonstrate that this is the case in two CNN models (a previously published model and a modification thereof) trained to estimate historical population-scaled recombination rate. Further, we observe that, when alignment sequence diversity is limited, including the associated positional information has mixed success in improving model performance. Our findings suggest that the previously observed inconsistency of CNN performance across recombination rates may be influenced by the amount of information in the corresponding alignments. CNN estimators perform poorly on alignments with low sequence diversity, i.e., when alignments have few segregating sites.

## Results

We investigate the limitations of CNNs in the context of recombination rate estimation. Population recombination rate, *ρ*, is defined as *ρ* = 4*Nr*, where *N* is population size and *r* is the crossover rate per base pair per meiosis. A previous study has shown that *ρ* can be estimated surprisingly well by CNNs, using as input only sequence data (Flagel et al., 2019). However, CNNs trained to estimate *ρ* are essentially black boxes: they take in population sequence data (paired genotype alignments and positional vectors derived from the original sequences) and output *ρ* values without any knowledge of the underlying biological processes. As a result, CNN performance is entirely dependent on the information that the network can learn from the labeled input data.

To understand the limitations of CNN *ρ* estimators, we first need to consider the data used in training and the information it contains. Genotype alignments are simplified representations of biological sequence data. Alignments can be best visualized as a matrix of 0s and 1s, where the rows correspond to sampled, phased chromosomes that have been aligned so that each column represents a site along the chromosome. For example, consider two independent, complete alignments of five sampled chromosomes sequenced across ten sites (Figure 1). All variable sites (also called segregating sites) have been encoded as 0s or 1s to indicate the presence of the reference (ancestral, 0) or alternate (derived, 1) allele. Sites that are conserved across all sampled chromosomes contain only 0s. Both alignments shown in Figure 1 have two segregating sites; however, the segregating sites vary in their location on the chromosome. This representation can be further reduced to two pieces of information: an alignment that only contains the segregating sites and a vector of locations of those segregating sites in the original chromosomes. The removal of conserved sites drastically reduces the computational power and memory required to work with large-scale genotype alignment data.

**Figure 1:**
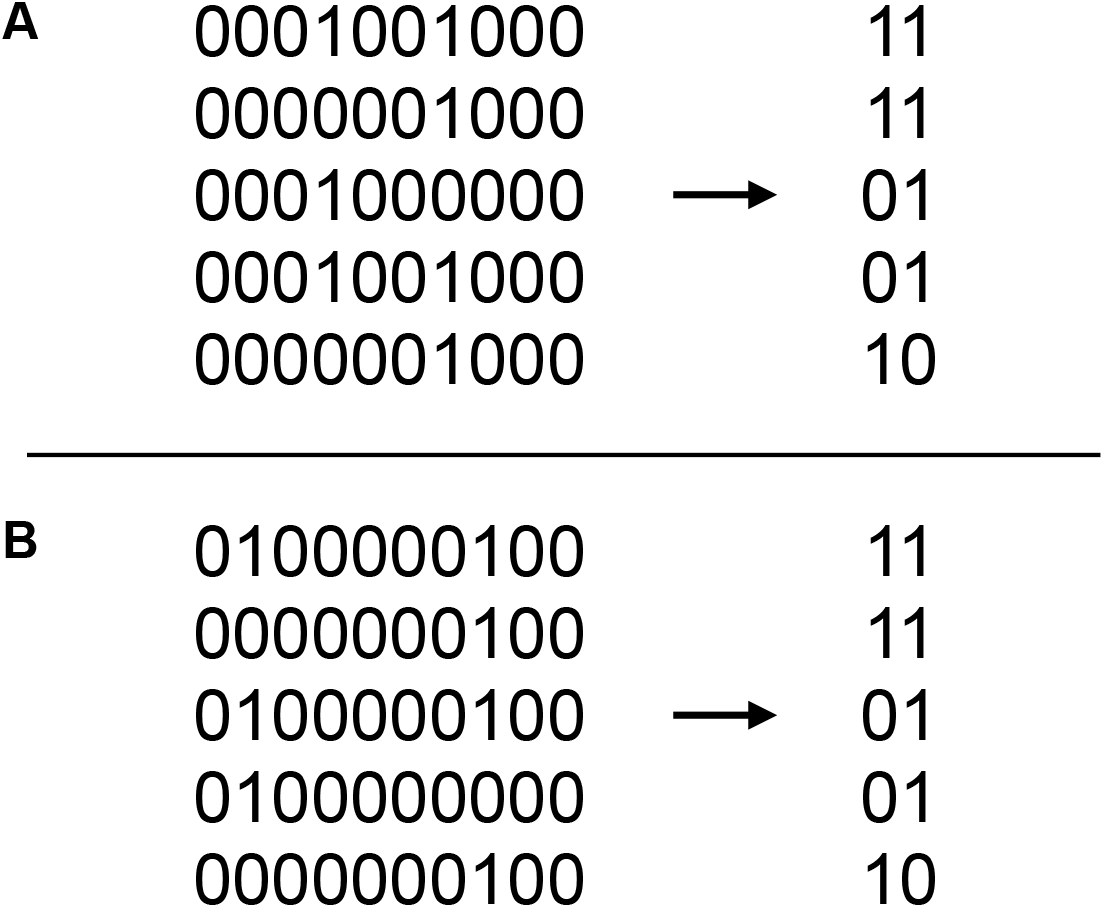
Duplicates are produced by removing conserved sites in sorted alignments. On the left, two independent, full sequence alignments are shown. Each row corresponds to a sampled chromosome, and each column represents a biallelic site along the chromosome. Alleles are encoded as 0s and 1s, where 0 corresponds to the ancestral/reference allele and 1 is a derived allele. The alignments considered in A and B both contain 2 variable (segregating) sites, with variation at sites 4 and 7 (panel A) or sites 2 and 8 (panel B). Genotype alignments used for model training include only segregating sites and have been reordered based on similarity. The result of this processing is two identical (duplicate) genotype alignments (shown on right). Positional vectors still vary between the identical alignments.

Genotype alignments are frequently subjected to an additional pre-processing step prior to being fed into recombination rate estimators; specifically, chromosomes are reordered within the alignment based on genetic similarity so that those most similar are sorted near each other. In the example of Figure 1, while we observe that the original alignments actually differ, the process of removing conserved sites and sorting results in two identical genotype alignments. We refer to this scenario as “duplicate” alignments. We expect that populations with little diversity (namely, those that produce alignments with few segregating sites) will produce duplicate alignments at considerable frequency.

While we know that some input parameter regimes may inherently produce data sets with limited diversity, it is not clear a priori which parameter settings are most problematic. We investigate this question by simulating populations with known historical recombination rates *ρ*, spanning several orders of magnitude. All data sets are generated with the ms coalescent simulation program, implemented in the msprime Python library (Hudson, 2002; Kelleher et al., 2016; Van Rossum and Drake, 2009). We study the sequence variation produced by different parameter regimes by systematically varying input parameters; i.e., we vary a single parameter at a time while holding others constant. Specifically, we consider a range of mutation rates (*µ*) and population sizes (*N*) in these simulations. We process our ms simulation output as one would for use by a CNN model or by a more conventional estimator such as LDhat (Flagel et al., 2019; McVean and Auton, 2007). Genotype alignments are separated from their positional information and, for CNN models, are subsequently sorted for similarity. Note that the order of sequences in an alignment is arbitrary, and thus sorting should be irrelevant. However, it helps with CNN training and convergence, as it reduces the input parameter space over which the network has to generalize its predictions.

Within each parameter regime (unique combination of *N* and *µ*), we do an all-by-all comparison of the 20,000 genotype alignments produced. We evaluate the genotype alignments for uniqueness before and after the additional pre-processing step of sorting. We find that certain parameter regimes produce duplicates at a substantial rate. Specifically, simulations with smaller population sizes *N* or mutation rates *µ* inherently have more duplicates (Figure 2). This effect is amplified when both mutation rate and population size are low.

**Figure 2:**
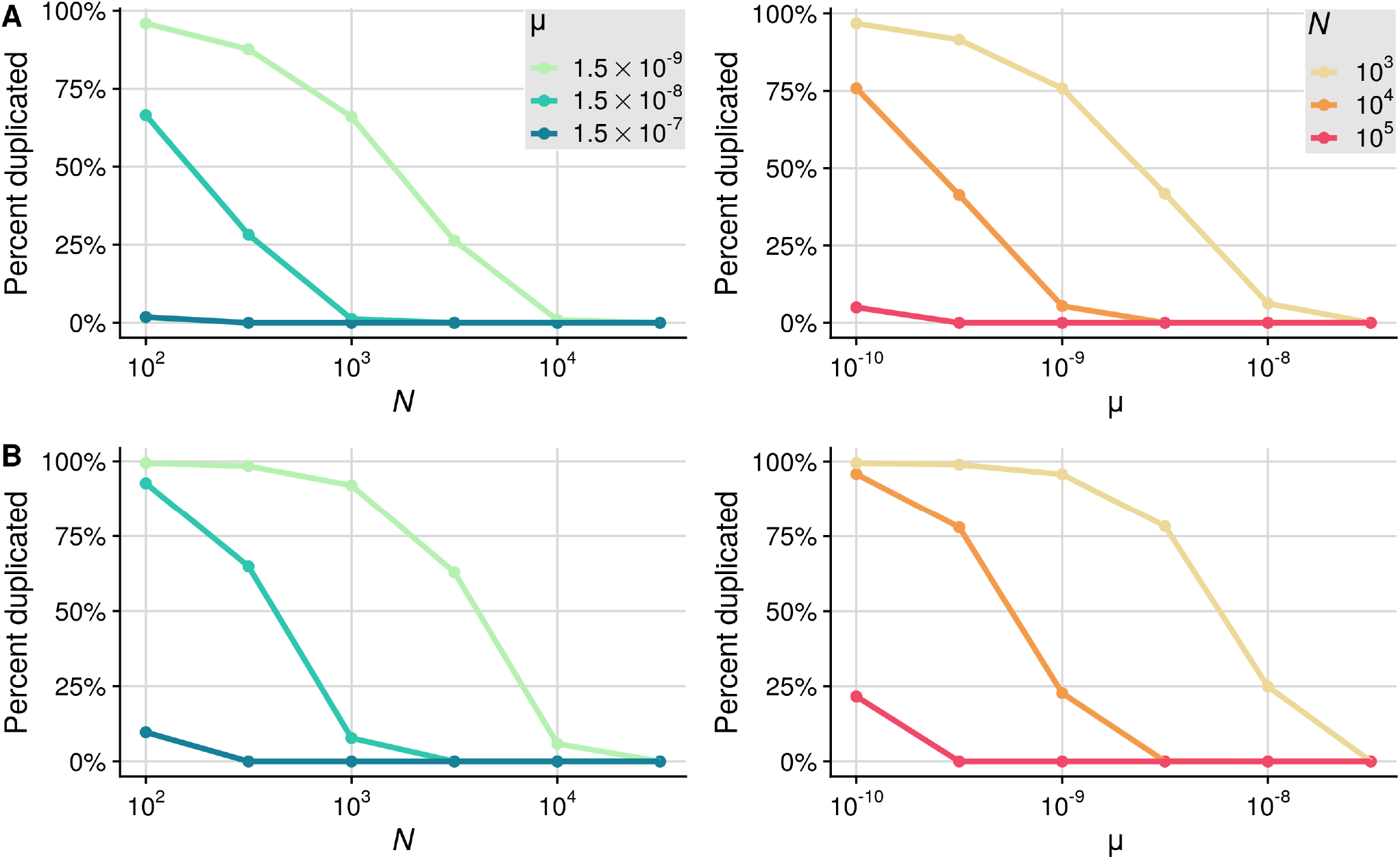
Percent of genotype alignments with at least one duplicate across simulation input parameter values. (A) Unsorted alignments. (B) Sorted alignments. Population size *N* and mutation rate *µ* vary across simulation sets, while all others input parameters are constant. Each point represents a set of 20,000 simulations in that parameter regime. We vary *N* logarithmically for 3 fixed *µ* values, and vice versa.

Flagel et al. (2019) found that sorting genotype alignments prior to training improves CNN performance. The sorting process allows the neural networks to more easily learn to estimate *ρ* because they do not have to generalize over many different, arbitrary orderings that inherently convey the same information and correspond to the same value of *ρ*. Because we are here focusing on CNNs for recombination rate inference, for the remainder of this work we consider exclusively the fully processed, sorted data sets. We note that sorting alignments for similarity increases the degree of duplication within a parameter regime by as much as 30% (Figure 2).

When we generate a training set for a CNN *ρ* estimator, our primary focus will be on the range of *ρ* values used to ensure it can accurately estimate all potential values. Here, we specifically generate data sets so that the simulations within each parameter regime span a comparable magnitude of *ρ* values, following the procedures introduced by Flagel et al. (2019). We visualize the relationship between the distribution of *ρ* values for the *N* and *µ* input values considered in the regimes shown in Figure 2 (left). By design, all regimes span a comparable magnitude of *ρ* values, with smaller *N* values corresponding to smaller *ρ* values (Figure 3). Because *ρ* does not depend on *µ*, the selected input values of *µ* do not impact its distribution. While these data sets seem interchangeable as training sets from the perspective of variation in *ρ*, they contain varying degrees of information. Parameter regimes associated with lower values of *µ* are more likely to contain redundant, duplicated alignments (Figure 2).

**Figure 3:**
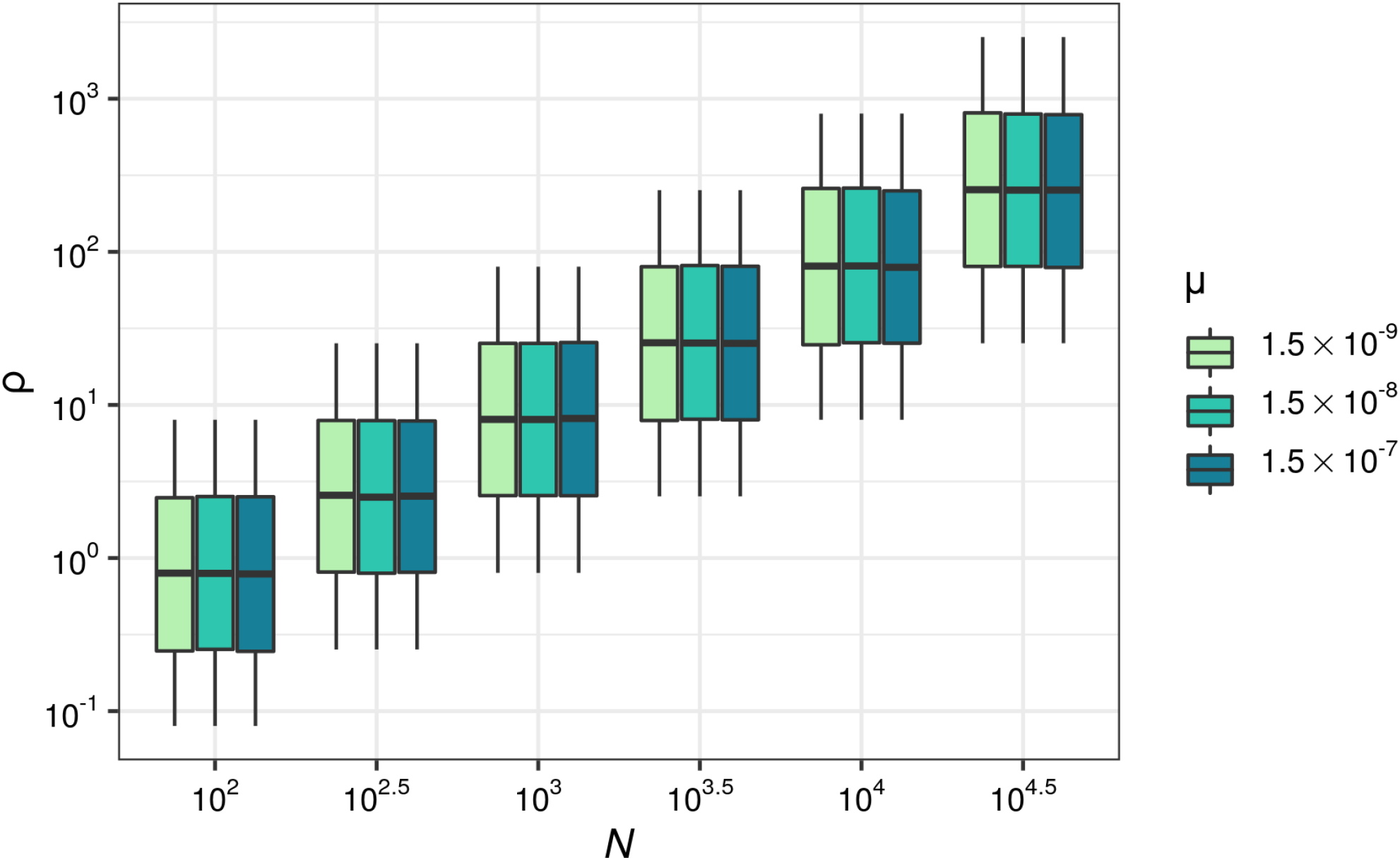
Distribution of *ρ* values used in simulations across associated parameter regimes. For the data sets considered in Figure 2, left, we show the *ρ* values against the population size *N* and mutation rate *µ* values used in simulations. Each unique *N* and *µ* combination represents a parameter regime with 20,000 independent simulations.

We have seen that duplicates arise in simulations with low population size and/or low mutation rate, and we have also seen that these parameter regimes are associated with lower values of *ρ* (Figure 3). Next, we ask where those duplicates fall across the range of *ρ* values within a simulation set. For every given population size *N* and mutation rate *µ*, we find that duplicate alignments are found to be evenly distributed across *ρ* (Figure 4A). We further examine characteristics of the resulting alignments to explain the occurrence of duplicates. We find that duplicates frequently occur in ms simulations when there are zero or one segregating (i.e., variable) sites in an alignment (Figure 4B). Duplicates are less frequent when the number of segregating sites exceeds one, but they still occur at non-negligible frequencies in alignments with two or three segregating sites (Figure 4B). Smaller population sizes inherently produce genotype alignments with relatively few segregating sites, and thus more duplicates. Similar results are found across all mutation rates and population sizes we considered (Figure S1). Importantly, the number of segregating sites is not a parameter in the simulations but rather an emergent property that displays substantial variation across replicates. Even if the expected number of segregating sites exceeds three, there may be a non-negligible fraction of simulations where the observed number of segregating sites is lower. As a consequence, a non-trival amount of duplicates may be observed in such parameter regimes (Figure S1, right-most panels).

**Figure 4:**
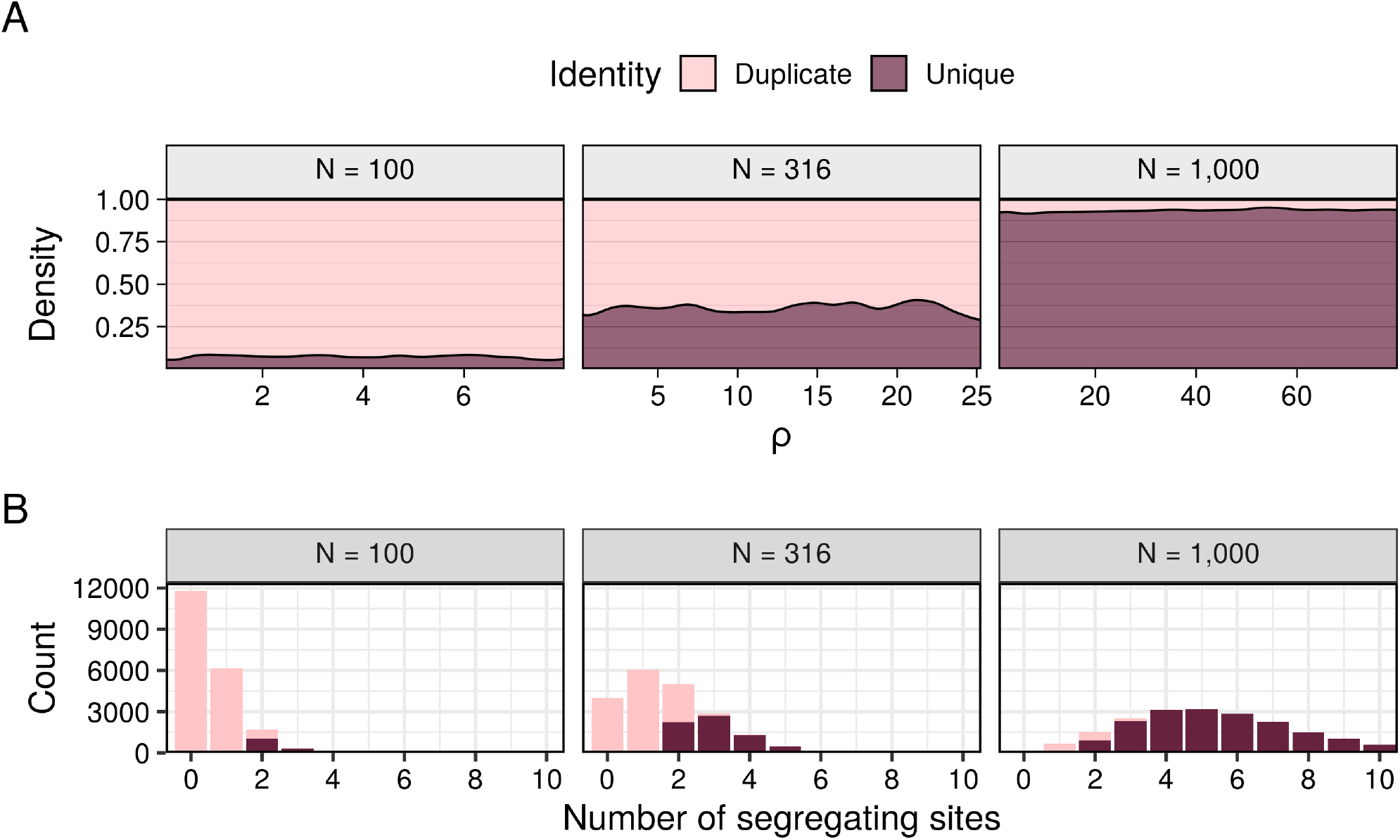
Distribution of *ρ* values and segregating sites for simulated alignments classified as either duplicate or unique in an all-by-all comparison of alignments within a given parameter regime. We consider a mutation rate of *µ* =1.5 × 10^−8^ and population sizes of *N* = 100, *N* = 316, and *N* = 1000 (left to right). Alignment identity (duplicate vs unique) is distinguished with color, where light purple represents alignments with at least one duplicate and dark purple for unique alignments. (A) The x axis has the range of *ρ* values generated in a regime, while the relative proportion of alignments based on identity (uniqueness) are depicted by the filled area. (B) The histograms show the distribution of alignments with between 0 and 10 segregating (or variable) sites.

We have observed that certain parameter regimes produce duplicate genotype alignments. Next, we investigate how these duplicates affect recombination rate inference by CNN. We consider two extreme cases by generating data sets with values on either end of the spectrum: one data set comes from a regime that produces many duplicates (referred to as “high duplicates” data set), while the other contains relatively few duplicates (referred to as “low duplicates” data set). The high duplicates data set includes alignments containing between 1 and 17 segregating sites, while the low duplicates data set includes alignments containing between 1 and 174 segregating sites (median number of segregating sites is 4 and 38, respectively). We use both data sets separately to train a CNN model to estimate recombination rate *ρ*, using the architecture described by Flagel et al. (2019). In each case, the model is trained for 18 epochs. This amount of training appears to achieve optimal performance, as can be seen from the relationship between the reported error on the training and validation sets (Figure S2, the validation error begins to diverge from the training error around epoch 18). This CNN, trained on paired alignment and positional data, performs relatively well on the low duplicates data set as reported by the performance metrics, *R*^2^ and RMSE, calculated on the test set (Figures 5 and S3, top left). The distance between the actual and estimated *ρ* values increases for smaller values of *ρ*, indicating that the CNN performs better for larger values of *ρ* on average. By contrast, the presence of duplicates at high abundance drastically decreases model performance for all *ρ* values (Figures 5 and S3, top right).

**Figure 5:**
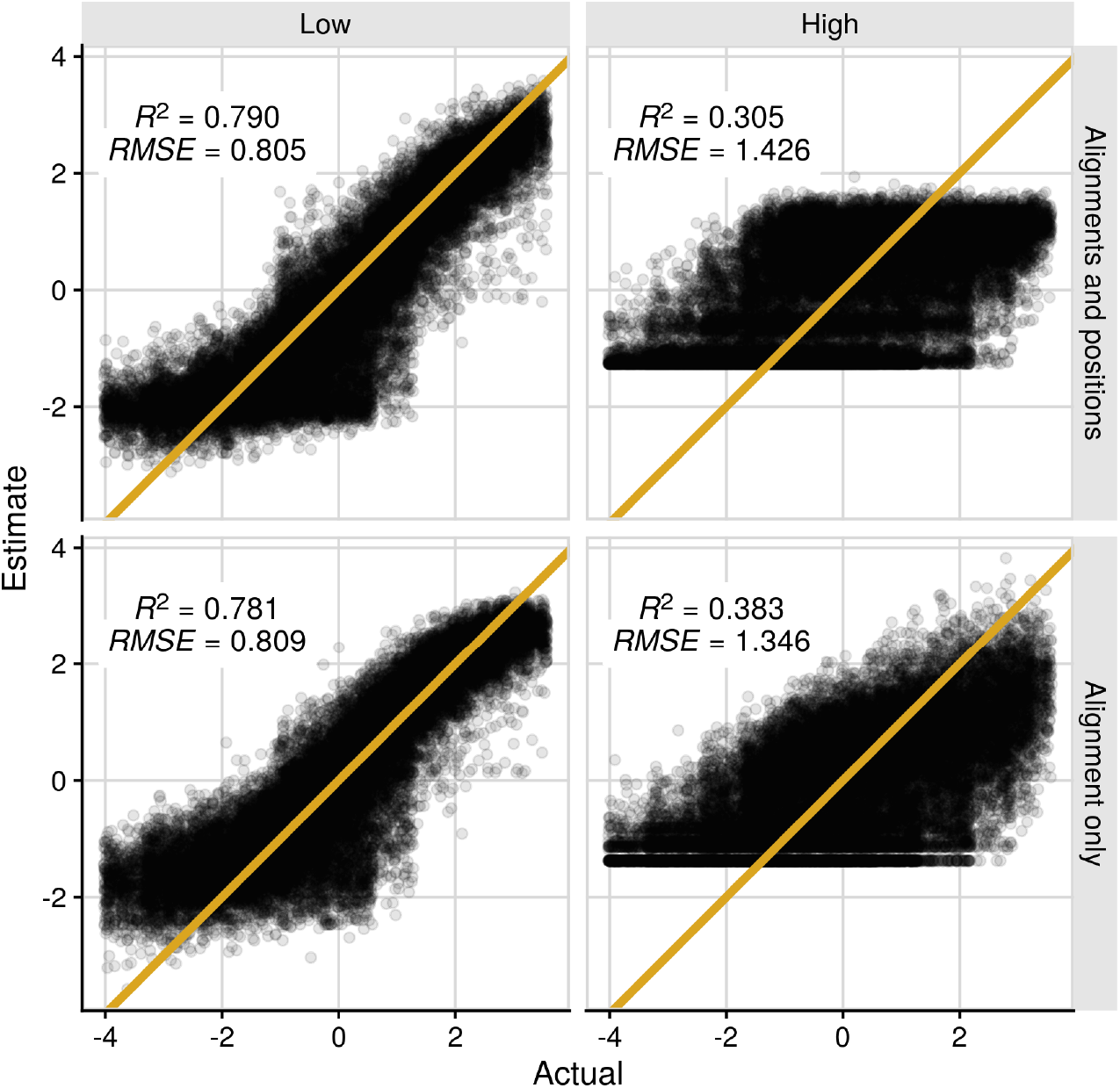
Performance of two distinct CNN architectures for *ρ* estimation trained on data sets with a low and high frequency of duplicate alignments. Plots show the log-transformed and mean-centered *ρ* values estimated by the trained CNN on each simulation in the test set against the actual (transformed and centered) *ρ* value used as input. The orange line references a 1-to-1 relationship that would be achieved if error was minimized to 0. All models are trained for 18 epochs before testing on a corresponding novel data set. The CNN model trained on genotype alignment and positional data (top, Flagel et al. (2019) architecture) achieves a RMSE of 0.805 on the low duplicate data set (left) and 1.426 on the high duplicate data set (right). With the removal of positional information and its corresponding branch, the alternate model (bottom) reaches an RMSE of 0.809 and 1.346, respectively, for the low and high duplicate data sets. Within each plot, the *R*^2^ and RMSE of the model on the designated test set are listed.

We further evaluate the impact of duplicates by training an alternate CNN. This model is a simplified version of the architecture presented by Flagel et al. (2019); rather than feeding in both genotype alignments and positional vectors separately into two network branches and later merging these branches into a combined network, we now utilize only the branch with alignment data. In other words, we entirely discard any positional information about where in the alignment the segregating sites occur. Naively, we would expect such a network to perform worse, as this positional information may both contain useful additional data about the recombination rate *ρ* and have the potential to disambiguate duplicates. (Two alignments that are identical except for the positional information will look entirely indistinguishable to this simplified network.) We find that the usefulness of the positional information is context- and model-dependent. For the high duplicates data set, removing the second branch and the positional vectors improves model performance in training (Figure S2, bottom) and testing (Figures 5 and S3, bottom left). By contrast, for the low duplicates data set, removal of positional information reduces model performance in training (Figure S2, bottom) and testing (Figures 5 and S3, bottom right). Importantly, discarding the positional information leads to a bigger increase in performance for high duplicates (7.8 percentage points increase in *R*^2^, from 30.5% to 38.3%) than it degrades performance for low duplicates (0.9 percentage points decrease in *R*^2^, from 79.0% to 78.1%). Thus, overall, it appears the CNN does not make efficient use of positional information. And in particular, in the parameter regimes where positional information would be the most critical, when there are many duplicates, the network actually performs better when it does not have access to this data.

## Discussion

We have characterized genotype alignments generated via coalescent simulations under a wide range of input parameters and corresponding recombination rates, and we have found that certain input parameter regimes limit the information contained within the alignments about the recombination rate. Unsurprisingly, information is limited when simulations are in a regime expected to produce low diversity; i.e., the simulated populations have a small population size *N* and/or a small mutation rate *µ*. Simulations in such parameter regimes generate alignments with relatively few segregating sites, which results in the same alignment being produced multiple times. We have further found that CNNs trained to infer recombination rate from genotype alignments are negatively impacted by the duplicates found in parameter regimes with low sequence diversity. One might assume that providing the paired positional data (locations of segregating sites) might provide enough distinction to overcome the information lost by duplicates. However, we have observed the opposite in a CNN that does not make use of this data: When duplicates are highly abundant, including positional information leads to a performance decline. Yet positional information increases model performance when duplicates occur at relatively low frequency.

Several groups have previously demonstrated the promising potential of deep learning inference to fill gaps left by traditional approaches to recombination rate estimation (Chan et al., 2018; Flagel et al., 2019; Adrion et al., 2020; Wang et al., 2021). We have here re-implemented a previously published CNN model (Flagel et al., 2019), and we have observed model performance similar to the reported performance when the training sets contained duplicates at low frequency. Notably, we have observed that the CNNs have the most difficulty estimating small *ρ* values, as previously reported. This region corresponds to parameter regimes that are most likely to contain duplicates, due to the correlation between *ρ* and *N*. We note that in Flagel et al. (2019)’s comparison of their CNN to a conventional method (LDhat), this was also the portion of the distribution where the CNN performed worse than LDhat (McVean and Auton, 2007). We speculate that the inferior performance of the CNN for small values of *ρ* could be caused, at least in part, by the presence of duplicate alignments in that part of the parameter regime.

Flagel et al. (2019) used a branched CNN with two separate inputs, genotype alignments and positional vectors. Here, we have additionally tested a variant architecture which ignores the positional vectors and estimates *ρ* using genotype alignment data alone. Our results suggest that the original approach, where genotype alignments and positional vectors are used as separate inputs, is vulnerable to performance declines in certain parameter regimes. Rather than helping with inference, the positional vectors seem to degrade network performance specifically when there are frequent duplicates, i.e., when positional information should be most relevant for accurate inference. This finding also suggests that the CNN estimators do not extract the maximum possible amount of information out of low-diversity alignments (since the positional data, which should be useful, is not making a difference), and it likely explains the prior observation that CNNs perform worse than traditional recombination rate estimators when *ρ* is small.

We have focused here on CNN estimators, but the implications of our findings are not limited to CNN models. The deep learning models developed for the task of recombination rate estimation have been benchmarked against conventional *ρ* estimators like LDhat and LDhelmet, and the boosted regression method FastEPRR (Flagel et al., 2019; Adrion et al., 2020; McVean et al., 2002; Chan et al., 2012; Gao et al., 2016). CNNs perform about as well as conventional methods overall, but offer performance benefits in the context of novel application and estimation when samples are limited. We expect that genotype alignments from regimes with limited diversity and information will suffer similar fates regardless of inference method; i.e., conventional methods will also see a reduction in performance with the inclusion of duplicates. The magnitude of this impact however may vary across methods. Inference methods based on explicit population genetics models may be better at taking advantage of positional information and thus outperform CNNs on alignments with few segregating sites.

We emphasize that the parameter regimes considered here have been selected to present extreme scenarios; training sets generated for realistic applications may not include as many duplicates as some of the regimes considered here. Nevertheless, our work highlights the importance of carefully evaluating the data sets used for model training. CNNs and other deep learning architectures learn to estimate recombination rate *ρ* from labeled training data alone, so their performance is dependent on the quality of the training data. It is imperative that information limits are identified and accounted for in model training.

We offer this analysis to emphasize an inherent limitation of genotype alignments. Not all parameter regimes have equivalent sequence diversity and thus information quality. As a result, caution should be used when applying deep learning inference to low diversity data sets. In the CNNs considered here, we have seen that variant architectures are able to extract more or less information from a given data set. Different architectures, specifically those that better leverage the information in paired alignments and positional vectors, might be better suited for sequences with low diversity. While multi-branch CNNs are commonly used, the best architectures and concatenation procedures have not yet been established (Khan et al., 2020; Yuan et al., 2020). By identifying a better representation for the data and an architecture that utilizes the full context of the data, further improvements to model inference ability could be possible (Alzubaidi et al., 2021).

One potential architecture that has already been successfully adopted for a closely related task is a recurrent neural network (RNN). CNNs are hierachical models frequently used in tasks like computer vision, while RNNs are sequence-based architectures developed for use in natural language processing or text prediction (LeCun et al., 1998; Elman, 1990; Dhruv and Naskar, 2020). Adrion et al. (2020) developed a RNN, trained on sequence data, to infer recombination landscapes by estimating local crossover rates (*r*) in sliding windows along the chromosomes. This neural network outperformed popular conventional and CNN *ρ* estimators (when *ρ* estimates were transformed to *r* values for comparison), especially in contexts where sample data was limited or missing (Flagel et al., 2019; McVean et al., 2002; Chan et al., 2012; Gao et al., 2016). Whether or not RNNs are similarly impacted by the inclusion of low-diversity alignments as the CNNs considered here remains unknown. A RNN *ρ* estimator might see performance improvements over a CNN estimator as has been observed for *r* estimation. As RNNs and CNNs represent and extract features from data in different manners, RNN architectures might be more suitable to leveraging the information in paired alignment and positional data.

## Methods

### Data generation

To train and test convolutional neural networks, we need labeled data. Here, that means genotype alignments sampled from populations with known historical recombination rates (*ρ*). We generate data sets with known *ρ* values via ms, a coalescent simulator which has been used for this purpose by others (Hudson, 2002; Flagel et al., 2019). Specifically, we run ms simulations with the mspms command line application in the msprime Python library (Van Rossum and Drake, 2009; Kelleher et al., 2016).

Each simulation takes a number of input arguments and produces a sample of chromosomes from a population evolved under a Wright-Fisher neutral model with constant population size. Program input arguments used here include: mutation rate (*θ*), recombination rate (*ρ*), number of loci (*L*), sample size, and number of replicates. For all simulations considered in our analysis, we vary mutation and recombination rates across simulations, while holding all other parameters constant. Following Flagel et al. (2019), we use a fixed number of loci undergoing recombination (*L* = 20 kb), maintain a sample size of 50 chromosomes per simulation, and generate only one replicate per function call. The population-scaled mutation parameter is thus defined as *θ* = 4*NµL*, where *N* is the population size and *µ* is the mutation rate per base pair per generation. Similarly, the population-scaled recombination rate is redefined as *ρ* = 4*NrL*, scaled across a constant genome size. Values of *r* are independently drawn from an exponential distribution with values spanning 10^−8^ to 10^−6^ for each simulation.

We generate data sets with variation in *θ* and *ρ* for two purposes: 1) to evaluate the prevalence of identical alignments across different parameter regimes, and 2) to evaluate the impact on model performance when training sets contain duplicates at either extreme. We create variation by altering the values of *µ* and *N* used in simulations. For our duplicates analysis, we consider various combinations of *µ* and *N*. In our first set of simulations, we consider 3 fixed values of *µ* (*µ* ∈ {1.5 × 10^−7^, 1.5 × 10^−8^, 1.5 × 10^−9^}) and vary *N* logarithmically (*N* ∈ {1.0 × 10^2^, 3.16 × 10^2^, 1.0 × 10^3^, 3.16 × 10^3^, 1.0 × 10^4^, 3.16 × 10^4^}). We additionally consider 3 fixed population sizes (*N* ∈ {10^3^, 10^4^, 10^5^}) and vary *µ* logarithmically (*µ* ∈ {1.0 × 10^−10^, 3.16 × 10^−10^, 1.0 × 10^−9^, 3.16 × 10^−9^, 1.0 × 10^−8^, 3.16 × 10^−8^}). For each combination of the considered *µ* and *N* values, 20, 000 independent simulations are generated, processed, and compared.

To generate data sets for model evaluation, we use two sets from either extreme. On one end, we have our high duplicate data set, which is generated under a parameter regime that guarantees a high proportion of alignments with exact matches. This set contains simulations with *µ* = 1.5 × 10^−9^ and *N* ∈ {1.0 × 10^3^, 2.0 × 10^3^, 5.0 × 10^3^, 1.0 × 10^4^, 1.5 × 10^4^, 2.0 × 10^4^}. We generate 20, 000 independent simulations per population size considered. We produce a low duplicate data set from the same range of population sizes, but push it towards the other end of the extreme by using a higher mutation rate (*µ* = 1.5 × 10^−8^). Again, 20, 000 simulations per *N* value are generated. The result is 120, 000 independent simulations for both the low and high duplicate data sets to be used in CNN training and testing.

### Data processing

From the standard ms output, we extract the following for each simulation: a genotype alignment, a variable position vector, and a recombination rate *ρ*. Genotype alignments are matrices of 0s and 1s, with 0s corresponding to the ancestral allele and 1s to the derived allele at a variable (segregating) site in the population. Position vectors contain the locations of all segregating sites in the population along the chromosome. For the purposes of CNN training, all input data types are processed with the standard procedures found to maximize model performance in prior work (Flagel et al., 2019). We prepare the data sets for all analyses as we would for deep learning. The specific pre-processing steps are described in the following paragraphs.

Across simulations, each alignment has a fixed number of chromosomes and a variable number of segregating sites ([50 × *n*]), and thus a variably-sized positional vector (*n*). All alignments and position vectors in a data set are required to be a consistent size for use in convolutional neural networks, so we pad all alignments and positional vectors to the maximum number of segregating sites found in that set. Padding is performed using the pad sequences function in the keras R package, which adds 0s and −1s to the right end of matrices and vectors (respectively) to enforce a consistent size across simulations (Allaire and Chollet, 2022).

Genotype alignments are additionally subjected to sorting and matrix operations prior to use in analysis and use in CNNs. The chromosomes, or rows, of each individual alignment are sorted for similarity based on their Manhattan distance (R Core Team, 2019). Chromosomes are reordered within their alignments accordingly. The sorting process introduces a logical order to alignments, which leads to improved training for CNNs (Flagel et al., 2019). Following sorting, alignments are transposed resulting in segregating sites as rows and chromosomes as columns, ([*n* × 50]). For the models considered here, this results in 1D convolutional filters being applied across segregating sites.

Within simulation sets, the distribution of *ρ* values is skewed towards 0. We balance data, and thus improve CNN training, by transforming the *ρ* values in training, validation, and test sets in the follow ways: all values within a set are log transformed then centered on the mean of the training set (by subtracting the mean from each value). The log mean centered *ρ* values are used in training, and as a result, the CNN models output *ρ* values on a similar scale. To directly compare model estimates with the standard range of *ρ* values, we must back-transform the estimates. We do so by adding the mean from the training data to each observation and then exponentially transform the values. Similar transformations are applied generally in deep learning and specifically in the context of this model (Flagel et al., 2019).

### Identifying duplicates

In our analysis, we quantify the information quality of different parameter regimes based on the degree of duplicated alignments found in the data set. When comparing individual simulations, we treat genotype alignments separate from their positional vectors. While this data is paired, it is treated separately by the network, being fed into two different branches. The convolutional layers learn based on alignment matrices alone, and thus we treat them as independent data to assess information quality in early network learning. The degree of duplication within a parameter regime (here, all 20, 000 simulations for each unqiue combination of *N* and *µ*) is determined through an all-by-all comparison of padded genotype alignments before and after sorting. If an individual alignment has at least one exact match within the set, it is labeled a “duplicate”. Alignments without an exact match are considered “unique”. We present the proportion of duplicates for each parameter regimes, and calculate it by dividing the total number of duplicates found by 20, 000, the total number of simulations considered.

### CNN model implementation, training, and evaluation

We re-implement the CNN architecture used by Flagel et al. (2019) to estimate *ρ*. The CNN has two branches, which independently consider the input data (alignments and positional vectors). It estimates the parameter label (*ρ*). Genotype alignments go through three 1D convolutional layers with ReLu activation, average pooling, and dropout. Positional vectors pass through a fully connected layer with dropout before being merged with the flattened output of the convolutional layers. The merged branches are then fed through two final, fully connected layers. In addition to this two branch model, we consider an alternate model that considers only genotype alignment information. The architecture follows that of Flagel et al. (2019) but excludes the positional vector branch with its data and fully connected layer; instead, the flattened output of the convolutional layers is passed to the final fully connected layers alone.

All models considered here utilize an Adam optimizer with a learning rate of 0.00001 and weight decay (or L2 regularization) parameter of 0.0001. Training was performed on CPUs using batch normalization (batch size = 32) and validation. We implement our CNNs via the R package torch (Falbel and Luraschi, 2022). Torch uses the functionality of the PyTorch natively in R by interfacing directly with its C++ library, libtorch (Falbel and Luraschi, 2022; Paszke et al., 2019).

We train each CNN independently for the 120,000 simulations in the low duplicate data set and 120,000 for the high duplicate data set (see subsection “Data generation” for details on parameter settings in simulated data sets). The respective sizes of padded alignments in these data sets are [174 × 50] and [27 × 50]. Following the standard convention, we split our data into training and validation sets (60% and 20%) and test sets (20%). Root-mean-square error (RMSE), which measures the error of the model in predicting *ρ*, is used as a metric for performance and model comparison (Twomey and Smith, 1995). RMSE is calculated between the model estimate of *ρ* and its label value (the ground truth). Training occurs over 18 epochs, which was found to be optimal for maximizing model performance while minimizing over-fitting to the training data by comparing the RMSE curves (Figure S2).

## Code

Data and code for this work are available at: https://github.com/mmjohn/genotype-alignment-information. All data analysis and figure production was performed in R version 3.6.1 (R Core Team, 2019), making extensive use of the tidyverse family of packages (Wickham et al., 2019). Additional functionality for data generation and prep come from a custom R package available at: https://github.com/mmjohn/popgencnn. All neural networks were implemented in the torch R package version 0.5.0 (Falbel and Luraschi, 2022).

## Acknowledgements

This work was supported in part by National Institutes of Health grant R01 GM088344. C.O.W. also received support from the Jane and Roland Blumberg Centennial Professorship in Molecular Evolution and the Dwight W. and Blanche Faye Reeder Centennial Fellowship in Systematic and Evolutionary Biology at The University of Texas at Austin. M.M.J. also received support from NIH training grant T32 LM012414-01A1. Computational resources for data generation and model training were provided by Texas Advanced Computing Center (TACC) and The Center for Biomedical Research Support (CBRS) at The University of Texas at Austin.

## Supplemental Material

**Figure S1:**
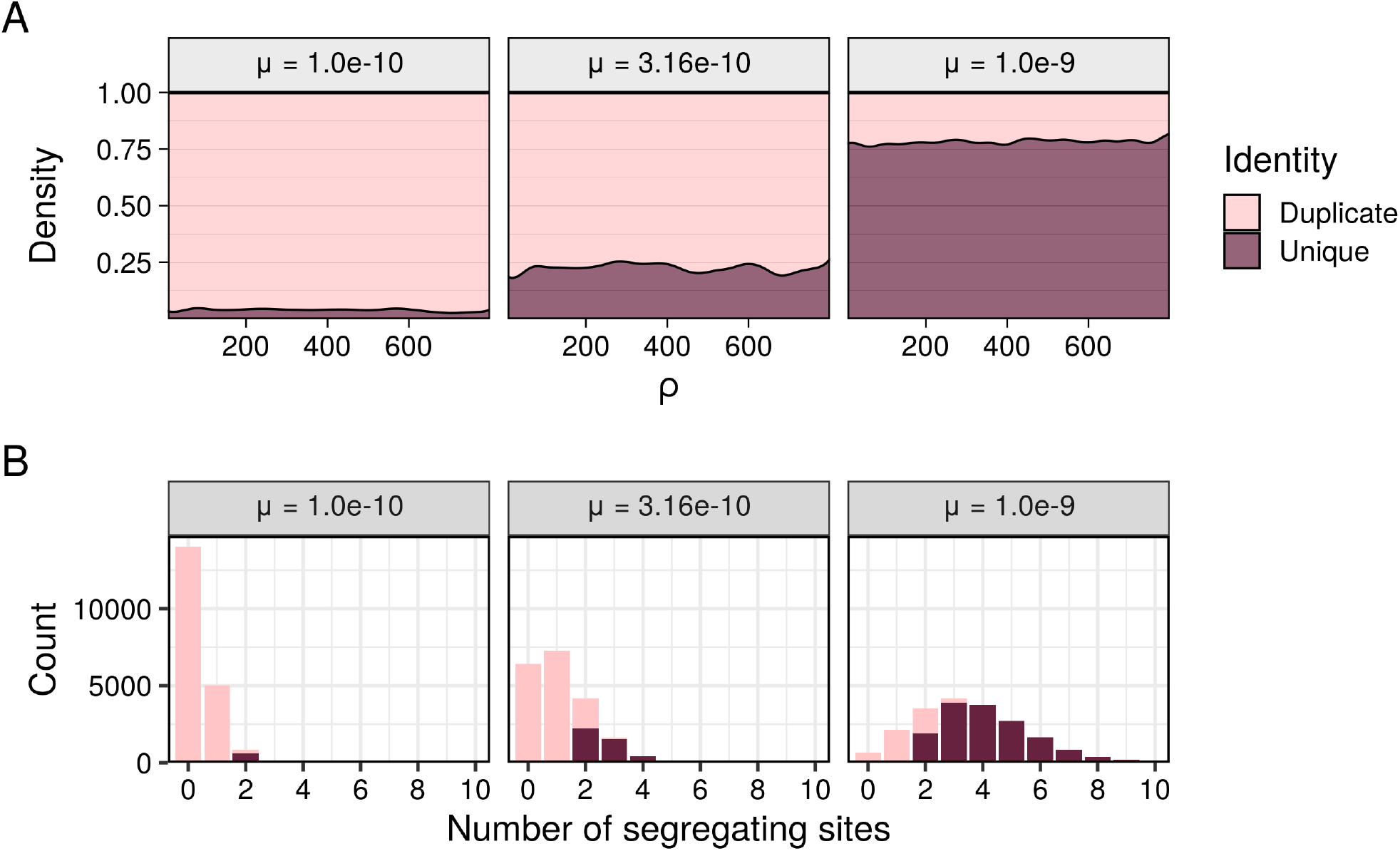
Distribution of *ρ* values and segregating sites for simulated alignments classified as either duplicate or unique in an all-by-all comparison of alignments within a given parameter regime. We consider a population size of *N* = 10^4^ and mutation rates of *µ* = 1.0 × 10^−10^, *µ* = 3.16 × 10^−10^, and *µ* = 1.0 × 10^−9^ (left to right). Alignment identity (duplicate vs unique) is distinguished with color, where light red represents alignments with at least one duplicate and dark red for unique alignments. (A) The x axis has the range of *ρ* values generated in a regime, while the relative proportion of alignments based on identity (uniqueness) are depicted by the filled area. (B) The histograms show the distribution of alignments with between 0 and 10 segregating (or variable) sites.

**Figure S2:**
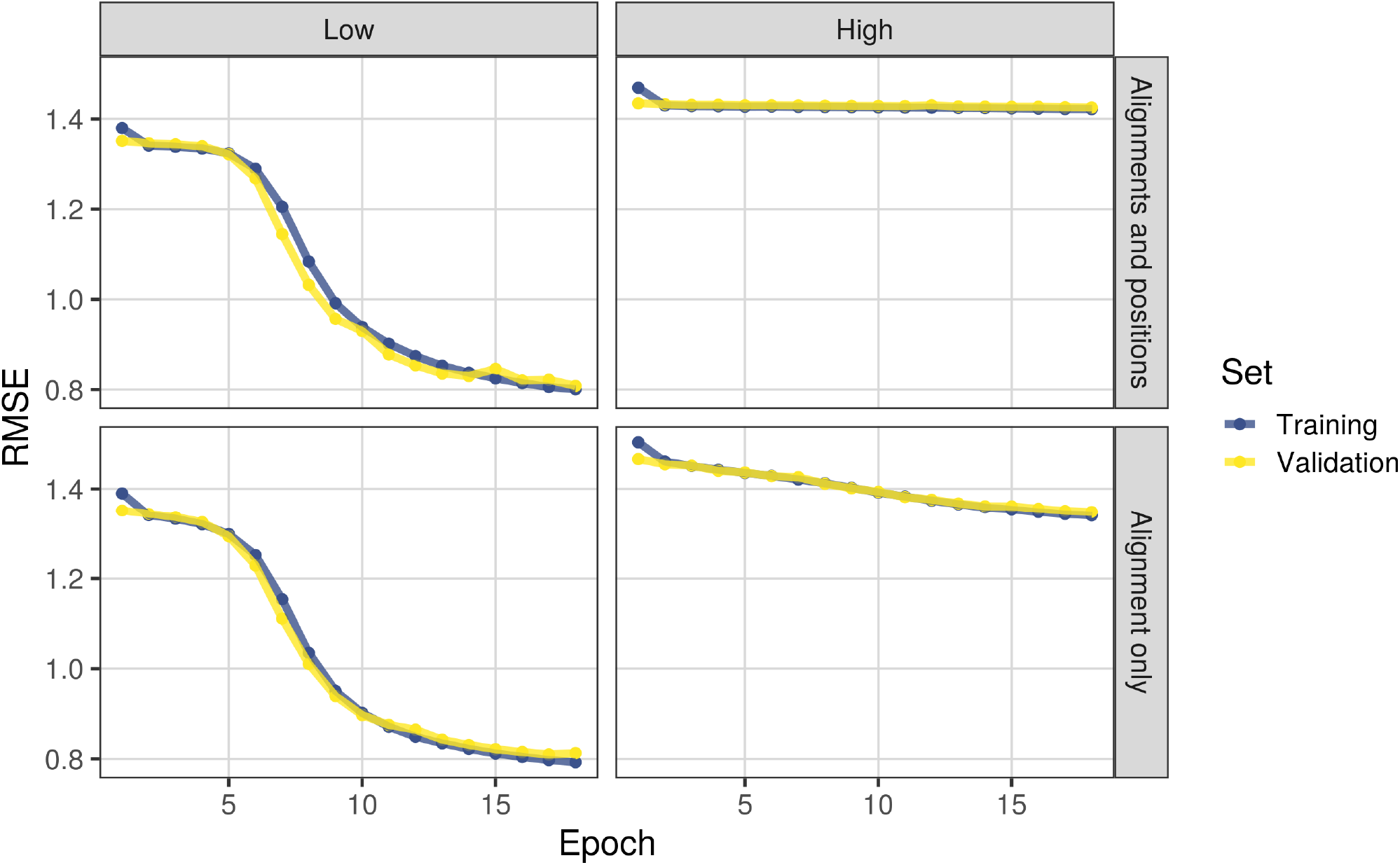
Training curves for a CNN *ρ* estimator and a variant architecture, using either genotype alignment data or alignment and positional data from each independent simulation, across 18 epochs. The estimator’s error (RMSE) is reported after each epoch on the training set (blue) and validation set (yellow). All models are trained on 72,000 simulations with a validation set containing 24,000 independent simulations. The low duplicate data set (left) has been simulated to contain relatively few duplicates, while the high duplicate data set (right) comes from a regime expected to produce duplicates frequently. Performance of each trained model on its respective test set is shown in Figures 5 and S3, presented as either log-transformed and mean-centered *ρ* values or population-scaled *ρ* values.

**Figure S3:**
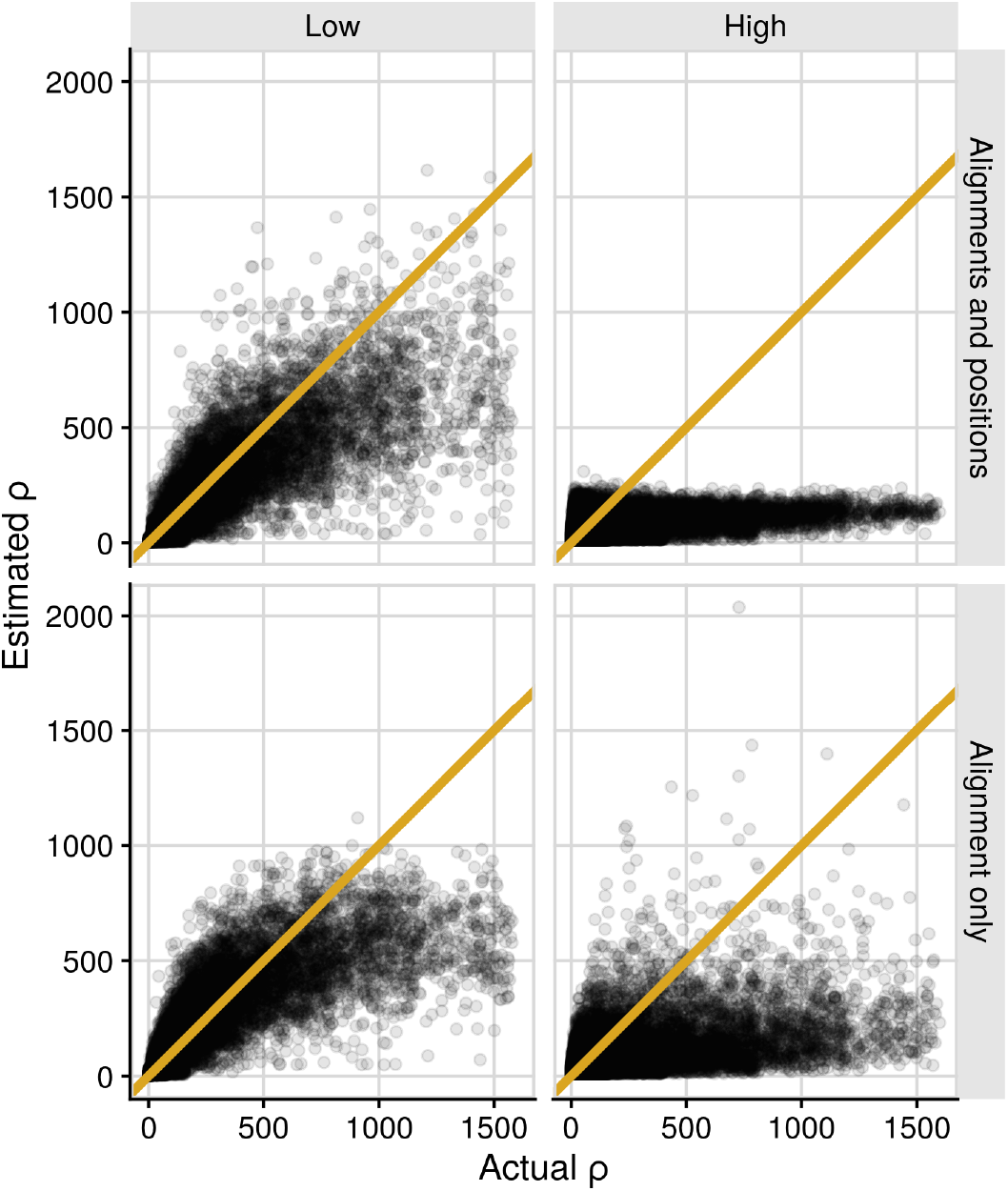
Performance of trained CNNs shown for untransformed *ρ* estimates. The orange line is included for reference to a perfect estimate (achieved if error is minimized to 0). The data points shown here are identical to those in Figure 5, except that *ρ* values now are not log-transformed or mean-centered.

